# Network Analysis and Human Single Cell Brain Transcriptomics Reveal Novel Aspects of Alpha-Synuclein (SNCA) Biology

**DOI:** 10.1101/2020.06.05.137166

**Authors:** Erin Teeple, Khushboo Jindal, Beril Kiragasi, Siddharth Annaldasula, Ann Byrne, Lilly Chai, Mahdiar Sadeghi, Can Kayatekin, Srinivas Shankara, Katherine W. Klinger, S. Pablo Sardi, Stephen L. Madden, Dinesh Kumar

**Affiliations:** Translational Sciences, Sanofi, Framingham, MA 01701, USA; Neurological and Rare Diseases Therapeutic Area, Sanofi, Framingham, MA 01701, USA

## Abstract

Alpha-synuclein (SNCA) aggregates are pathological hallmarks of synucleinopathies, neurodegenerative disorders including Parkinson’s Disease (PD) and Lewy Body Dementia (LBD). Functional networks are not yet well-characterized for SNCA by CNS cell type. We investigated cell-specific differences in SNCA expression using Allen Brain Database single-nucleus RNA-seq data from human Middle Temporal Gyrus (MTG, 15,928 nuclei) and Anterior Cingulate Cortex (ACC, 7,258 nuclei). Weighted gene co-expression analysis (WGCNA) and hierarchical clustering identified a conserved SNCA co-expression module. Module genes were highly conserved (p < 10^−10^) and most highly expressed in excitatory neurons versus inhibitory neurons and other glial cells. SNCA co-expression module genes from ACC and MTG regions were then used to construct a protein-protein interaction (PPI) network, with SNCA empirically top hub. Genes in the SNCA PPI network were compared with genes nearest single nucleotide polymorphisms linked with PD risk in genome-wide association studies. 16 genes in our PPI network are nearest genes to PD risk loci (p < 0.0006) and 55 genes map within 100kb. Selected SNCA PPI network genes nearest PD risk loci were disrupted by CRISPR knock out gene editing for validation of network functional significance; disruption of STK39, GBA, and MBNL2 resulted in significantly elevated intracellular *SNCA* expression.

## INTRODUCTION

Nervous system alpha-synuclein (SNCA) aggregates are characteristic findings in Parkinson’s Disease (PD), Lewy Body Dementia (LBD), Multiple System Atrophy (MSA), and Pure Autonomic Failure (PAF), a group of disorders collectively referred to as the synucleinopathies (1). While the synucleinopathies share the presence of pathological SNCA aggregates associated with dysfunction and death of nervous system cell populations, these disorders differ with respect to the anatomic locations of SNCA pathology, the cell populations primarily impacted, and even in the intracellular locations where aggregations are found (1). For example, neuronal Lewy bodies and Lewy neurites are observed in LBD and PD, but with differences in the involved brain regions (1, 2). In further contrast, SNCA aggregates are present as oligodendroglial cytoplasmic inclusions in MSA (1, 3) and as peripheral nervous system neuronal cytoplasmic inclusions in PAF (1, 4). SNCA is also not observed exclusively as a neuropathological finding in the synucleinopathies: up to 50% of Alzheimer’s Disease (AD) cases also exhibit SNCA aggregates (5). Whether and to what extent SNCA aggregations occur as cause, consequence, or both among these different neurodegenerative disorders remains incompletely understood. The specific processes by which different degenerative cellular neuropathological processes share SNCA aggregations as a common finding and how SNCA participates in the initiation and progression of each of these conditions remain areas of ongoing research.

SNCA is a small, 140-amino acid protein consisting of an N-terminal region containing imperfect KTKEGV repeats, a hydrophobic middle domain, and a highly negatively charged C-terminus(6). It is a member of the synuclein protein family, and in neuronal cells has been found to localize both to the nuclear and cytoplasmic compartments (7–9). In neurons, SNCA can be found in high concentrations at synaptic terminals associated with the synaptic vesicles(7, 10). SNCA has been found to interact with presynaptic proteins to function as a chaperone to maintain pre-synaptic soluble N-ethylmaleimide-sensitive factor attachment protein receptor (SNARE)-complex assembly during neurotransmitter release (11, 12). Yet the activity of SNCA does not appear to be isolated to synaptic transmission. SNCA has also been found to reduce DNA methylation by interacting with and sequestering maintenance DNA methyltransferase (Dnmt1) in the cytoplasm (7, 13), and to interact directly with histone proteins and DNA in the nuclei of cultured cells, with effects on transcription and cellular toxicity responses (9, 14). These interactions support more complex functional and regulatory roles for SNCA. Intracellular SNCA balance depends on the autophagy-lysosome pathway (ALP) for aggregation and degradation of SNCA, with ALP inhibition shown to reduce intracellular SNCA aggregation, increase extracellular SNCA secretion, and provoke cellular inflammatory responses (15, 16).

The polygenetic inheritance patterns and measurable contributions of environmental and epigenetic factors to synucleinopathy risk provide further support for nuanced participation of SNCA at the intersection of multiple intracellular pathways. Genetic variants associated with synucleinopathy risk include not only SNCA overexpression and structural modification mutations (17), but also variants linked to mitochondrial dysfunction, lysosomal storage disorders, oxidative stress responses, and alterations in potassium channel function (18–20). Susceptibility genes shown to increase PD susceptibility include GBA, PARK2, PARK7, PINK1, and LRRK2, which connect PD risk with multiple pathways(21–24). Understanding the ways in which SNCA participates in and connects such diverse cellular processes as are represented among syncleinopathy-linked genes and protein expression variants is a necessary step for better understanding and to identify potential treatments and biomarkers for SNCA-related disorders.

While heterogeneities in cell types and brain regions involved in synucleinopathies present challenges for understanding this protein, recent advancements in single-cell and single-nucleus sequencing have enabled new analyses of cell-specific variations in gene expression. Here, using full transcriptomic RNA-seq data from the Allen Brain Database for samples from the human Middle Temporal Gyrus (MTG, 15,928 nuclei) and Anterior Cingulate Cortex (ACC, 7,258 nuclei) (25), we investigated pathways and cell-specific differences in cortical SNCA expression. Single-nuclei RNA-Seq analysis is a relatively recently developed method which may be applied to identify heterogenous subpopulations and characterize transcriptomic differences among cells in single tissue samples (26).

Among the cell groupings identified in our analysis, we observed conserved patterns of differential neuronal SNCA expression in ACC and MTG regions, with high SNCA expression observed in excitatory (glutamergic; GLUT) neurons and low expression in inhibitory (gabaergic; GABA) neurons. A protein-protein interaction network was then constructed for genes in the identified conserved SNCA co-expression module based on known protein-protein interaction pathways. Among the 1495 genes identified as being part of this SNCA PPI network, many were either nearest genes to highly significant PD risk loci identified in genome-wide association studies (20), or within ±100 kb of highly significant PD risk loci. CRISPR knockouts of selected genes tended to increase alpha synuclein levels, further confirming the functional significance of our SNCA PPI network.

## MATERIAL AND METHODS

### Data Sources

The Allen Brain Cell Types Database contains high quality full-transcript single nuclei, SMART-Seq data with a median gene capture of up to 9000 genes (25). We downloaded and processed single-cell RNA-seq data from this database for human middle temporal gyrus (MTG) (15,928 nuclei) and anterior cingulate cortex samples (ACC) (7,283 nuclei). These ACC and MTG data sets are publicly available and were generated from samples of frozen human brain specimens from 8 healthy human tissue donors (age range: 24-66 years; MTG and ACC samples from same donors). From the downloaded data files for each anatomic region, intron and exon gene-level read counts were combined to produce single files for analysis (MTG or ACC). Gene expression data were then filtered to include only human protein-coding genes, excluding those on the X and Y chromosomes (27).

### SNCA Module Detection Using Weighted Gene Correlation Network Analysis (WGCNA)

Weighted gene correlation analysis (WGCNA) was performed in R using the WGCNA package (28) (Fig. 1). For each sample location (MTG and ACC), a gene-gene co-expression matrix was first generated from the normalized cell-gene arrays extracted from the Seurat data objects. As per WGCNA protocol, a scale free-topology fit index was plotted as a function of potential values for the soft thresholding power, and a soft threshold power of 8 was selected empirically for optimal transformation of each co-expression similarity matrix into a topological overlap matrix (TOM). The TOM matrix reflects relationships of topological similarity between genes. The dissimilarity matrix (1 – TOM) is used to represent dissimilarity, and by convention, this dissimilarity matrix is then used to cluster groups of genes into co-expression modules (28). This ensures that co-expressed genes are similar at both the expression and network topology levels. Hierarchical clustering and dynamic cutting were then applied to identify modules of co-expressed genes. The dynamic tree cutting algorithm (deep split = 2) was used to detect gene modules, which are defined as clusters of densely interconnected genes in a co-expression network. The module containing the SNCA gene (the cluster of genes co-expressed with SNCA) was identified for ACC and MTG. The ACC and MTG SNCA modules were then overlapped to find common genes between them and to identify a robust module with conserved expression across sample locations for further downstream use. Statistical comparison of module overlaps was performed in R using the Hypergeometric test function phyper(), where q = number of genes overlapping between MTG and ACC; m = gene MTG module size; n = total number of genes (estimated to be 20,000) – MTG genes to find number of non-MTG genes; and k = gene ACC module size.

**Figure 1.**
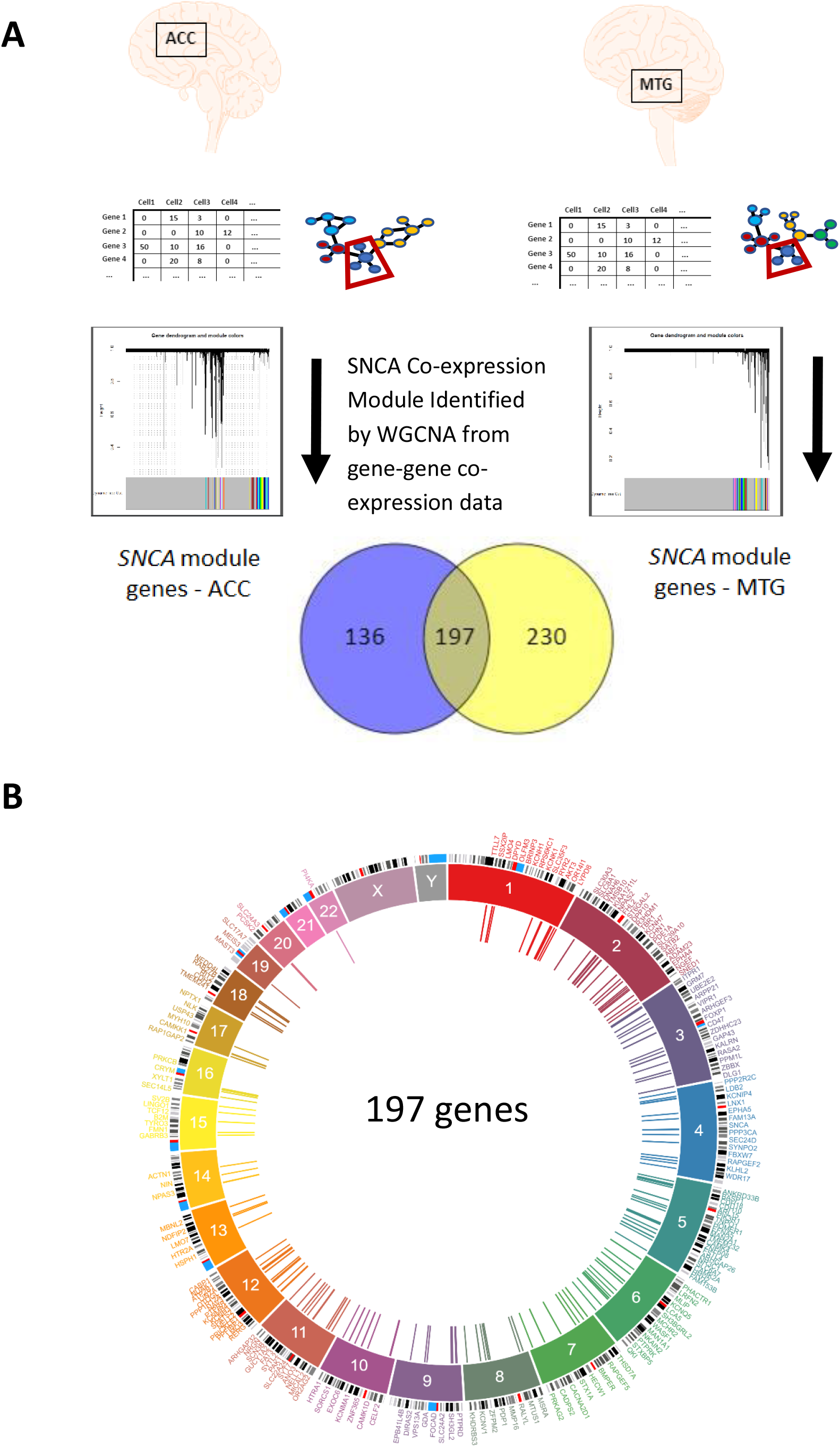
SNCA Co-expression module conserved across sample locations identified by WGCNA hierarchical clustering. (A) Genes belonging to SNCA-containing cluster are those co-expressed with SNCA. (B) SNCA co-expression module gene beds mapped to human genome (inner circle: module genes).

### SNCA Module Hub Gene Identification by Protein-Protein Interaction (PPI) Network Analysis

Genes identified by WGCNA as being part of the SNCA co-expression module conserved across MTG and ACC locations could be correlated with SNCA by random chance rather than having a true functional interaction with SNCA. A protein-protein interaction (PPI) network was thus created for genes identified as belonging to the conserved SNCA co-expression module in order to integrate the identified robust module with information on known protein-protein interactions and to determine hub genes/proteins within this network (29). NetworkAnalyst and the STRING database were used to generate and visualize this PPI network and to identify additional interacting proteins connected within the network generated for the robust SNCA co-expression module genes (29).

### Single Nuclei RNA-Seq Analysis of Human MTG and ACC Tissue Samples

Single-cell RNA-Seq analyses for MTG and ACC were performed in R using the Seurat package, version 3.0 (Fig. 2) (26). A standard data pre-processing workflow was applied using cutoffs 200 < nFeatureRNA < 9500 and percent.mt < 0.01 for MTG and 200 < nFeatureRNA < 8500 and percent.mt < 0.01 for ACC, with cutoffs selected based on initial QC plots. Data were normalized using global-scale log normalization, scaling by a factor of 10,000 for each data set. Identification of highly variable features, linear dimension reduction by PCA transformation, and cell clustering were performed using standard Seurat package workflows with PCA dimensions optimized to achieve separations by cell type markers (n = 17). Cell type annotations for each cluster were determined by cluster marker visualizations and marker distributions quantified using feature and violin plots. SNCA expression within each cluster was examined by labelling nuclei by expression level of the SNCA gene. Differential SNCA expression was then compared by examining cluster staining on TSNE plots and by using dot plots to compare differences in mean expression by cluster and proportion of cells within each cluster expressing SNCA.

**Figure 2.**
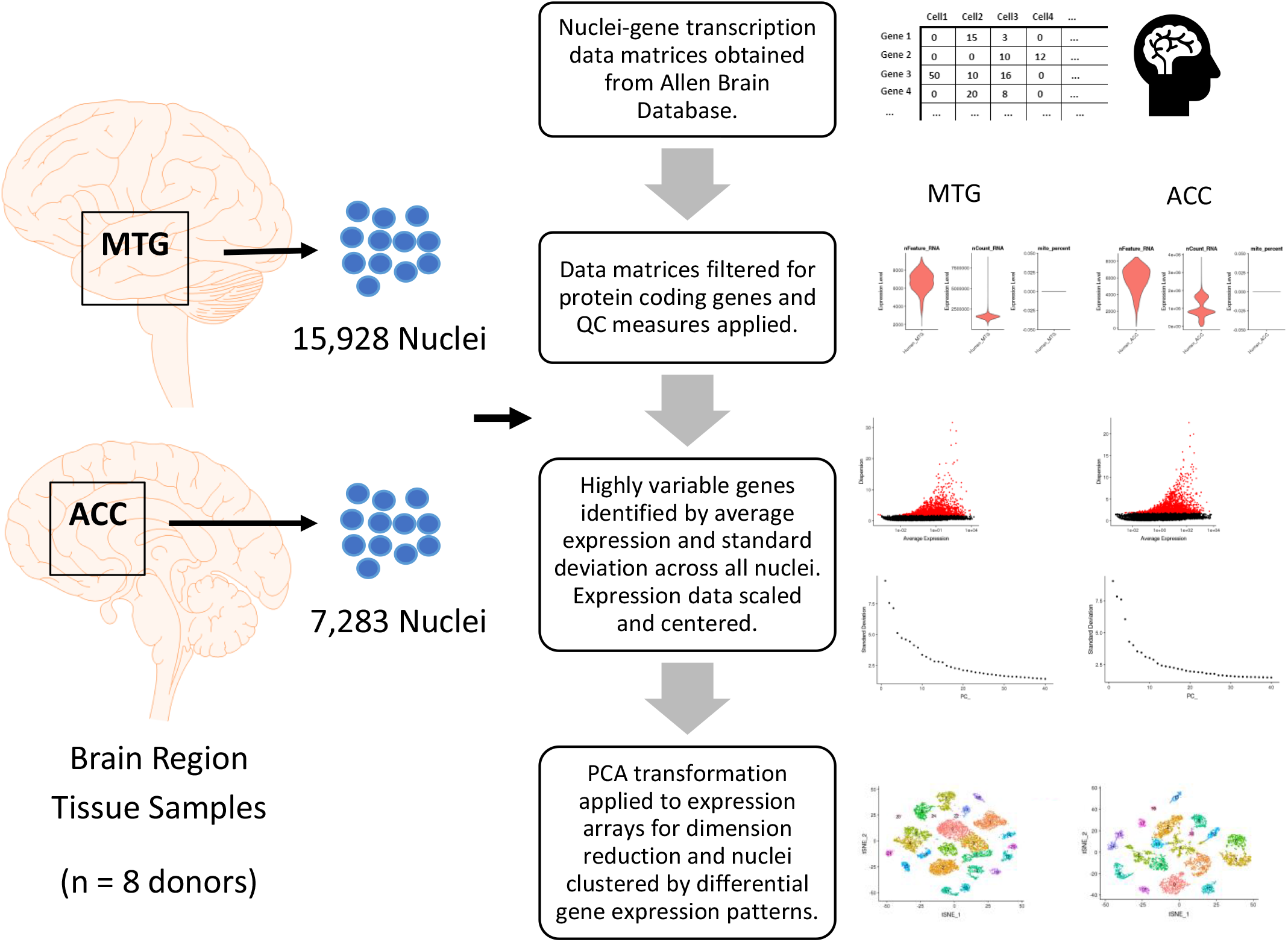
Single-cell RNA-Seq analysis of human MTG and ACC samples. Schematic overview of data pre-processing, quality control (QC), and variable gene and PCA dimension selection workflow.

### SNCA Module Gene Ontology and Comparison with PD Genome-Wide Association Studies

Genes identified from the SNCA module and PPI interaction network were also compared with a list of genes identified as being nearest genes to single nucleotide polymorphisms (SNPs) linked with significantly increased risk for Parkinson’s Disease in genome-wide association studies and with a larger set of genes mapping to regions within ±100 kb of highly significant SNPs (20). To identify genes within 100 kb of highly significant variants, variants reported in the summary statistics tables for Nalls and colleagues’ 2019 meta-analysis of genome wide association studies for PD (PD GWAS) were searched using a p-value cutoff for PD association of less than 10^−5^ (n = 5806; ‘nallsEtAl2019_excluding23andMe_allVariants.tab’) (20). Human genes within ±100 kb of these SNPs were identified using Ensembl and UCSC genome annotations for Hg19/Gr37 and the R package ‘BioMart’ (27, 30), yielding a set of 709 unique genes. We then compared the sets of nearest genes and genes in ±100 kb beds with the 197-gene union set of the ACC and MTG modules and the ACC and MTG co-expression module PPI network. Statistical tests for overlap enrichment were performed in R using the hypergeometric test. Overlap visualizations for each of these steps were generated using Venny (31). The Ingenuity Pathway Analysis (IPA) tool (Qiagen) was used to identify networks and processes involving genes for the robust SNCA module obtained from the intersection of the ACC and MTG locations and the intersection of this module with genes represented in the PPI network analysis and comparison with genes adjacent to PD GWAS loci (32).

### Alpha Synuclein Levels in CRISPR Knock Out Cell Models

#### Mammalian Cell Culture and Cell Line Generation

In order to examine how SNCA module/PD risk gene disruptions might impact SNCA protein expression, CRISPR-Cas9 knock outs were created for the genes SNCA, STK39, GBA, MBNL2, DLG2, NSF, FCGR2A, RIT2, NECAB2, and SH3GL2. The human neuroblastoma cell line SH-SY5Y was obtained from Sigma (Cat no. 94030304) at passage number 5 and used for generation of all other CRISPR-Cas9 edited cell lines. Cells were kept in a humidified incubator at 37 °C and 5% CO2. All cells were maintained in DMEM high glucose, + L-glut, -sodium pyruvate media (Cat no. 11-965-092; Gibco by Life Technologies) supplemented with 10% heat-inactive FBS (Cat no.A3840001; Thermo Fisher Scientific) and 1% Pen/Strep (Cat no. 15-140-122; Gibco by Life Technologies). Cas9-expressing cell pool was generated by lentiviral delivery of Cas9 (Cat no. <u>A32064</u>; Thermo Fisher Scientific) followed by selection with Blasticidin (5ug/ml) and was tested for Cas9 expression by Western Blot. CRISPR knock out cell lines were generated by lentiviral delivery of 4 gRNAs targeting each gene (see supplemental table for gRNA sequences) followed by selection with puromycin (5 ug/ml). These gRNA sets were custom generated by Thermo Fisher Scientific. The control cell line was generated by lentiviral delivery of (Cat no. A32062; Thermo Fisher Scientific) gRNA sequence with no sequence homology to any region of the human genome, followed by selection with Puromycin (5 ug/ml). Successfully transduced cell pools were used for further analysis.

#### Immunofluorescence Labeling, Confocal Microscopy and Image Analysis

11 different CRISPR knock out cell lines were seeded at 30,000 cell density per well into 96-black well, clear bottom Corning high content imaging plates in fresh growth media (selected genes plus scrambled control). Upon 24 hr incubation, cells were fixed in 4% paraformaldehyde (Cat no. 28906; Thermo Fisher Scientific) for 10 min, then blocked and permeabilized in 1% BSA (Cat no. 05470; Sigma) + 0.05% Saponin (Cat no. 47036; Sigma) in PBS supplemented with Ca^2+^ and Mg^2+^. Washes were performed 3x for 5 min. Fixed cells were incubated overnight at 4 °C with primary mouse monoclonal anti-alpha synuclein antibody (Cat no. 610787; BD Biosciences; 1:200) and for 2 hr with secondary goat anti-mouse Alexa Fluor Plus 488 antibody (Cat no. A32723; Thermo Fisher Scientific; 1:1000). Cells were then incubated overnight at 4 °C with PE-conjugated LAMP1 CD107a clone H4A3 antibody (Cat no. 527993; Molecular Probes; 1:20). After 3× washes, cells were treated with 2 ug/ml Hoechst 33342 (Cat no. H3570; Sigma) in PBS for 10 min and imaged with a confocal microscope (Opera Phenix). Images were acquired using 63× magnification 1.15 NA water objective with 9 fields per well and a z-stack of 4 planes, bright field included. Mean fluorescence intensities were quantified for each cell using Harmony software. Non-specific staining by the secondary antibody was controlled by incubation without primary mouse monoclonal anti-alpha synuclein antibody. Fluorescence intensity measurements were compared using one-way ANOVA followed by Dunnett’s multiple comparison test.

## RESULTS

### SNCA co-expression module is conserved across human MTG and ACC regions

WGCNA and hierarchical clustering applied to normalized gene count transcription matrices for all nuclei identified clusters of genes co-expressed with SNCA for MTG (n = 427 genes) and ACC (n = 333 genes) samples. From these SNCA co-expression clusters for MTG and ACC, we identified a statistically significant intersection set of 197 genes (hypergeometric p-value = 5.3e-247), which consisted of genes in the SNCA-containing co-expression clusters for both MTG and ACC. This intersection set of SNCA-co-expressed genes comprises a robust co-expression module for cortical SNCA (Fig. 1). The robust and statistically significant overlap between ACC and MTG suggests the conservation of the SNCA co-expression module across different regions of the cortex.

### Single Nuclei RNA-Seq data from Human MTG and ACC have representation of all major cell types

Nuclei-gene matrices for MTG and ACC samples from the Allen Brain Database were filtered and processed using the Seurat package workflow to annotate cell type for nuclei in these gene count transcription matrices (Fig. 2) (26, 33). For each location (ACC and MTG), nuclei were clustered based on highly variable genes using an unbiased graph-based clustering approach. Cell types for these unbiased clusters were annotated using broad expression markers for excitatory/glutamergic neurons (GLUT; SLC17A); inhibitory/GABAergic neurons (GABA; GAD2), astrocytes (ASTRO; GFAP), oligodendrocytes (OD; MOG), oligodendrocyte precursor cells (OPC; PDGFRA), and microglia (MG; CSF1R) (Figure 3). These annotations confirmed the presence of all major cell types in both ACC and MTG with a higher percentage of excitatory neurons observed in MTG (67.3% clustered nuclei) versus ACC (55.6% clustered nuclei). Neuronal nuclei comprised the majority of sample nuclei for both ACC and MTG samples (Fig. 3; Table 1).

**Table 1.** Nuclei counts by cell annotation; GLUT: glutamergic neurons; GABA: GABAergic neurons, OD: oligodendrocytes; ASTRO: astrocytes; OPC: oligodendrocyte precursor cells; MG: microglia.

| **Cell Cluster** | **GLUT** | **GABA** | **OD** | **ASTRO** | **OPC** | **MG** |
| --- | --- | --- | --- | --- | --- | --- |
| **MTG** | 10675 | 4164 | 360 | 344 | 244 | 84 |
| ***proportion*** | *0.673* | *0.262* | *0.023* | *0.022* | *0.015* | *0.005* |
| **ACC** | 3980 | 2282 | 211 | 341 | 169 | 179 |
| ***proportion*** | *0.556* | *0.319* | *0.029* | *0.048* | *0.024* | *0.025* |

**Figure 3.**
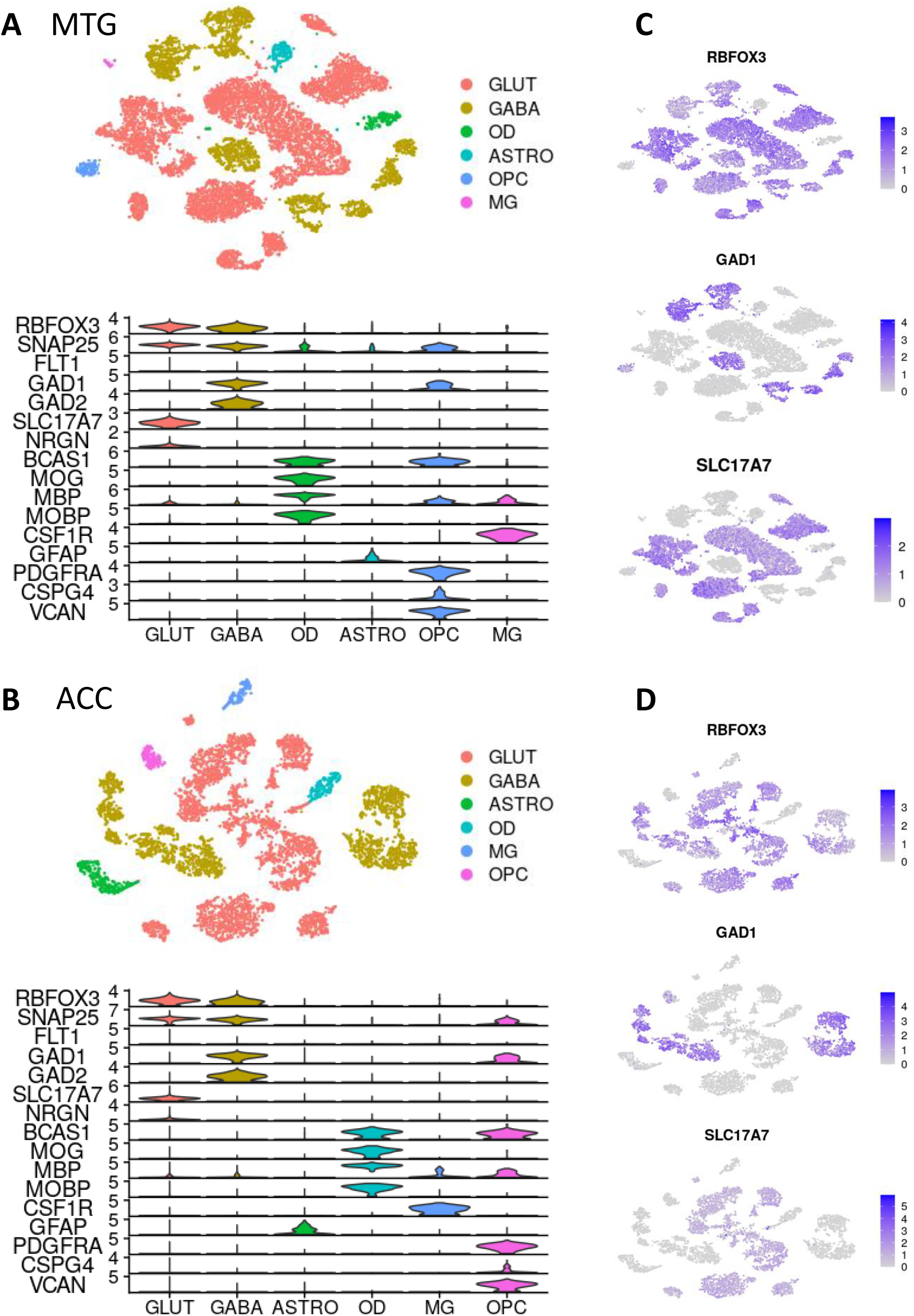
**Cluster annotations using cell type markers** for (A) MTG and (B) ACC. Feature plots for show marker expression delineated by clusters for (C) MTG and (D) ACC.

### SNCA and SNCA co-expression module genes have enriched expression in excitatory (glutamatergic) neurons

We then sought whether SNCA and the conserved 197 genes in the co-expression module were differentially expressed by cell type in ACC and MTG. To explore this question, we plotted heatmaps and feature plots from the annotated, normalized single cell data (Fig. 4). Genes in the SNCA co-expression module were observed to be most highly expressed in excitatory neurons in comparison to inhibitory neurons and other cortical cell types for both ACC and MTG samples. Expression levels for genes in the 197-gene co-expression module for the annotated cell type clusters are presented for MTG (Fig. 4A) and ACC (Fig. 4D). Alongside the observed increased expression of SNCA co-expression module genes, greater expression of SNCA itself was also observed in excitatory neurons in comparison to inhibitory neurons and all other non-neuronal cell types (MTG: Fig. 4B, C; ACC: Fig. 4E, F).

**Figure 4.**
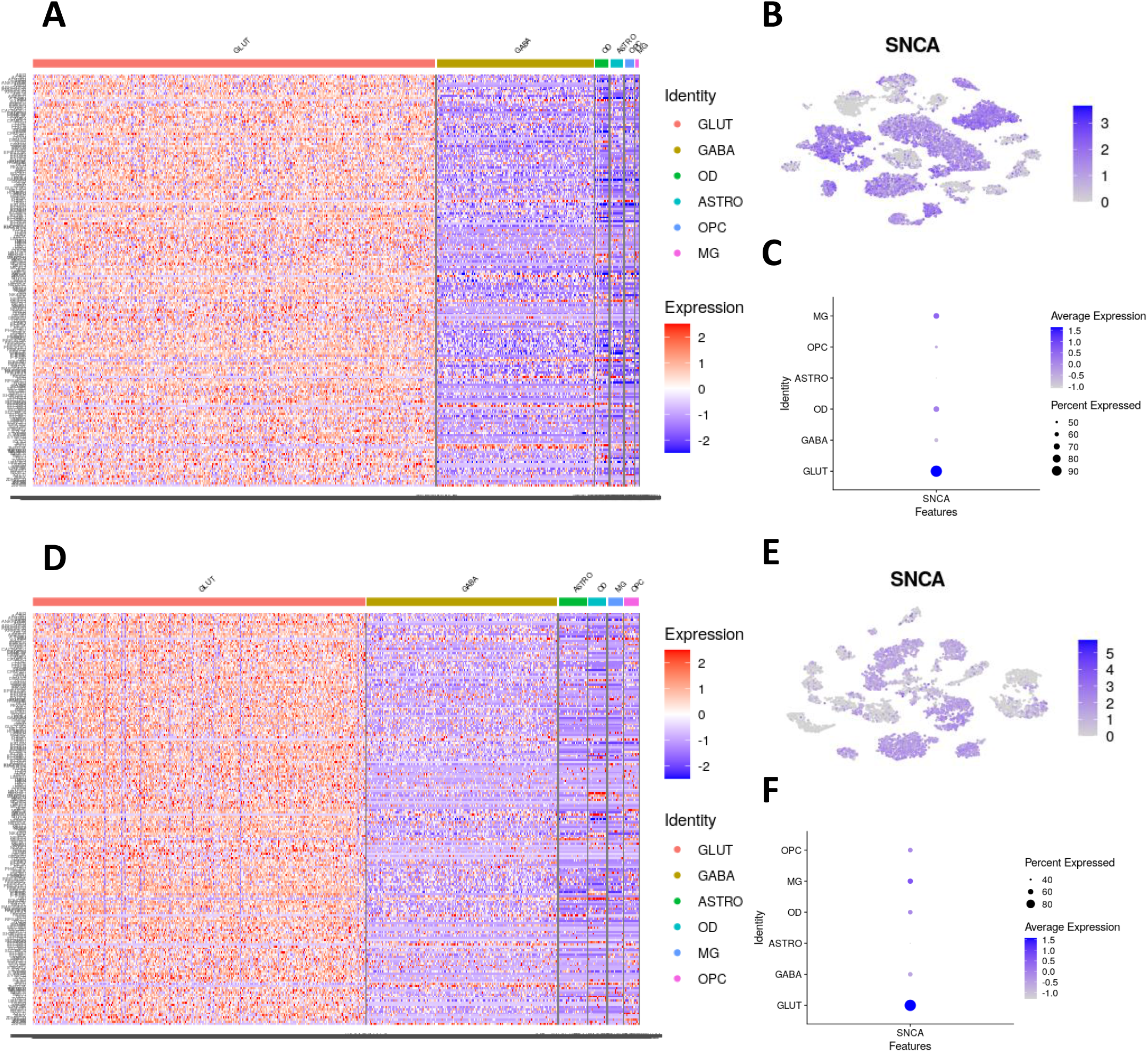
SNCA module gene expression and SNCA expression by cell type for ACC and MTG. Excitatory neurons are observed to have greater expression of SNCA co-expression module genes(MTG-A,B; ACC-D,E) and SNCA (MTG-C; ACC-F) compared with inhibitory neurons and non-neuronal cell types.

### SNCA co-expression module genes are implicated in synaptic biology and dopamine processing

Synaptogenesis, nitric oxide, calcium and ephrin A signaling, as well as dopamine feedback pathways were found to be the top pathways significantly enriched for among genes co-expressed with SNCA in ACC and MTG when this gene set was analyzed using IPA (Fig. 5A). Pathway analysis for the 197-gene conserved SNCA co-expression module also identified statistically significant enrichment for molecular targets of the upstream regulators Levodopa, Histone Deacetylase (HDAC1), cAMP Responsive Element Binding Protein 1 (CREB1), and SNCA, among others (Fig. 5B). Gene ontology (GO) analysis further identifies cellular localization, synaptic transmission, and ion transport as among functions enriched for in the set of 197 conserved SNCA-co-expressed genes (Fig. 5C) (34, 35). The identified pathways and regulators found to be enriched for among SNCA-co-expressed genes are central to synaptic biology and dopamine processing, supporting a role for SNCA as a multifunctional protein acting at the intersection of multiple cellular pathways.

**Figure 5.**
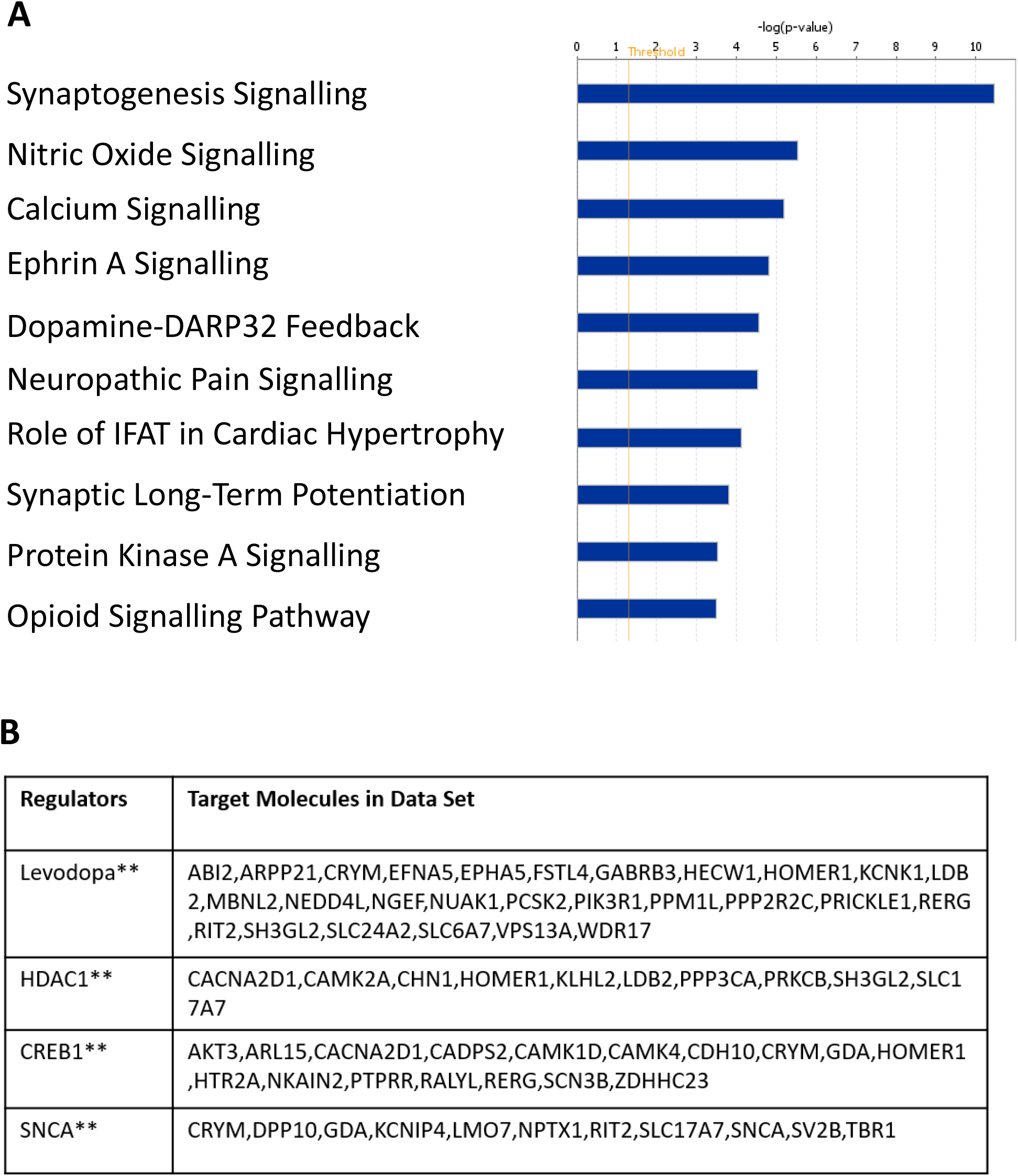
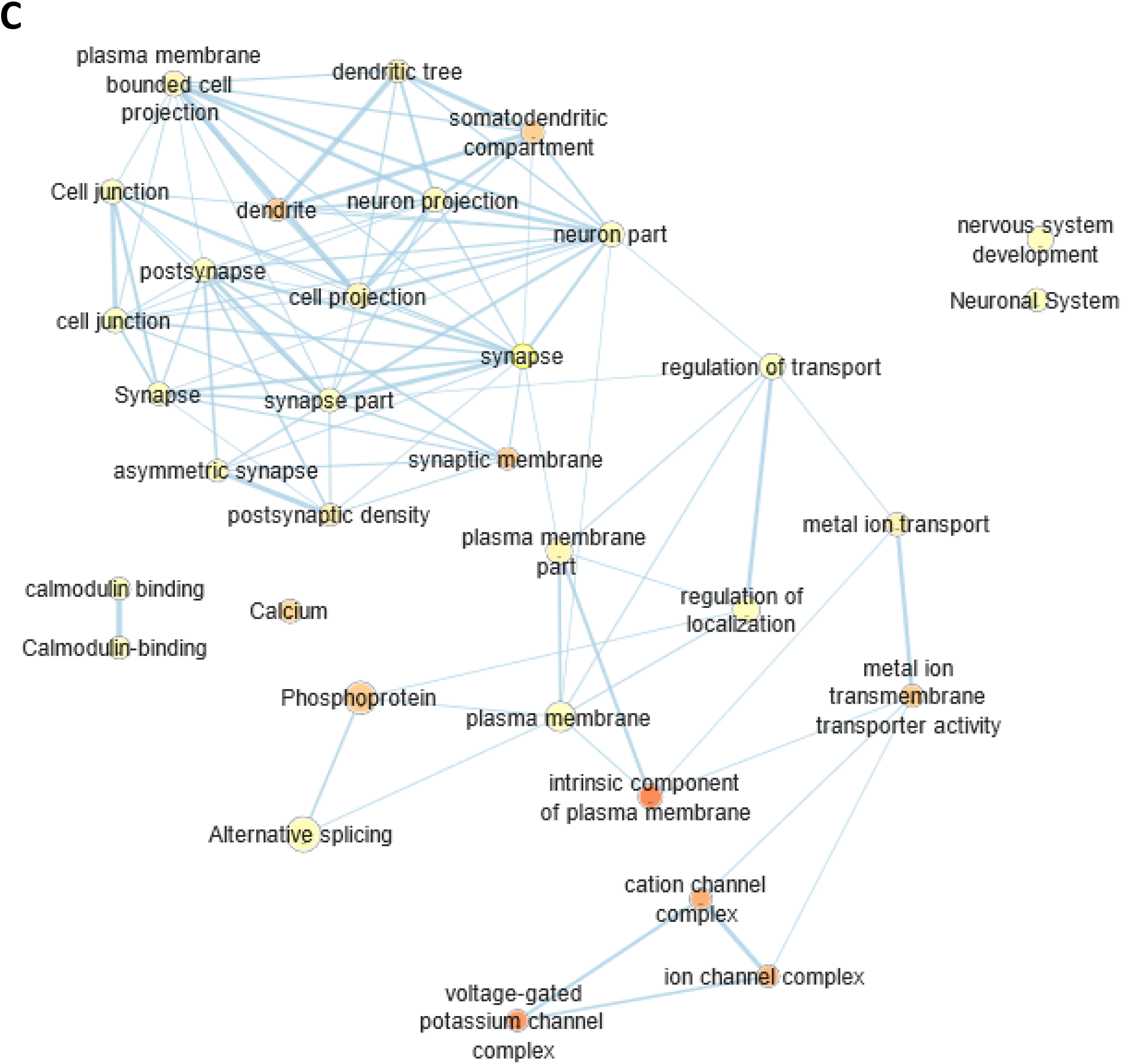
**Ingenuity Pathway Analysis for 197-gene conserved SNCA co-expression module** for nervous system tissue excluding cancer pathways (A) Top canonical pathways and (B) Top regulator molecules enriched for among module genes (**overlap p < 10-4). (C) EnrichmentMap Analysis reveals functional enrichment for cellular localization, synaptic transmission, and ion transport (q < 10^−5^; darker red coloration indicates more significant pathway enrichment).

To further explore whether SNCA and its 197-gene conserved co-expression module might be altered in the synucleinopathy Parkinson’s Disease (PD), differential expression of these 197 SNCA co-expression module genes was queried for in archival studies comparing PD case and control samples obtained for human cortex, putamen, substantia nigra, and dopaminergic neuron tissues. Records for this search were extracted from OmicSoft DiseaseLand Database (release HumanDisease_B37 20171220_v7) using an adjusted p-value cutoff of 0.05. Heatmap comparison of these search results reveals numerous significant differences in PD case versus control expression of SNCA co-expression module genes as well as tissue-specific variations in SNCA module gene expression (Fig. 7A). Consistent with our findings in MTG and ACC, frontal cortex samples have a number of genes with significantly increased expression for PD samples versus controls, and in addition, dopaminergic neurons are also found to have increased expression of these genes for PD versus Control samples (Fig. 7A; tissue sample type annotation indicated by x-axis color bar).

### SNCA is the major hub protein in the substantia nigra-specific PPI network derived from co-expression module genes

To obtain further insight into the functional pathways connecting SNCA and its co-expressed genes, the 197 genes in the conserved co-expression module were then used to generate a protein-protein interaction network using NetworkAnalyst (29), restricting our network model to substantia nigra-specific interactions. Network analysis yielded a network of 1495 proteins connecting the genes in our identified module via known protein-protein interactions (Fig. 6A). SNCA was empirically found to be the top hub of this network by both degree and betweenness criteria (Fig. 6B), further establishing SNCA as a central participant in interacting neural pathways. Interaction pathways within this PPI network potentially offer opportunities for targeted intervention to modulate SNCA expression or biomarker identification to monitor SNCA expression pathways in the CNS. We also used this network to identify other top hubs (Fig.6B); top PPI network hubs ordered by degree (highest to lowest) are the following: SNCA, FHL2, LNX1, ACTN1, PIK3R1, NR3C1, SH3GL2, NEDD4L, QKI, and FBXW7.

**Figure 6.**
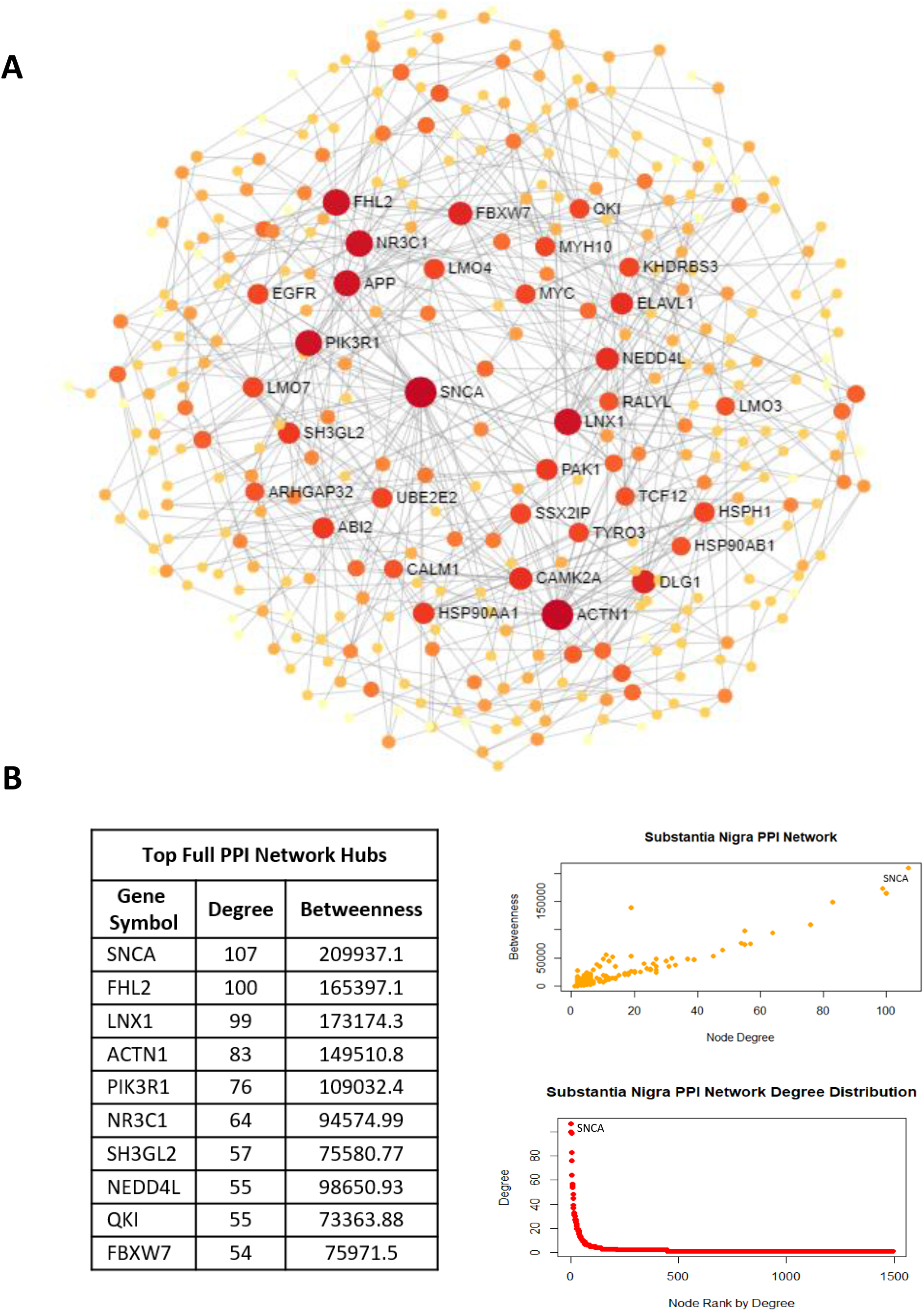
Protein-protein interaction network generated for the genes in the conserved *SNCA* co-expression module. (**A**) Minimally connected protein-protein interaction network for genes co-expressed with *SNCA* is shown with degree represented by node size and color (greater degree: greater size and darker red). (**B**) *SNCA* is the top hub in this network by both degree and betweenness.

**Figure 7.**
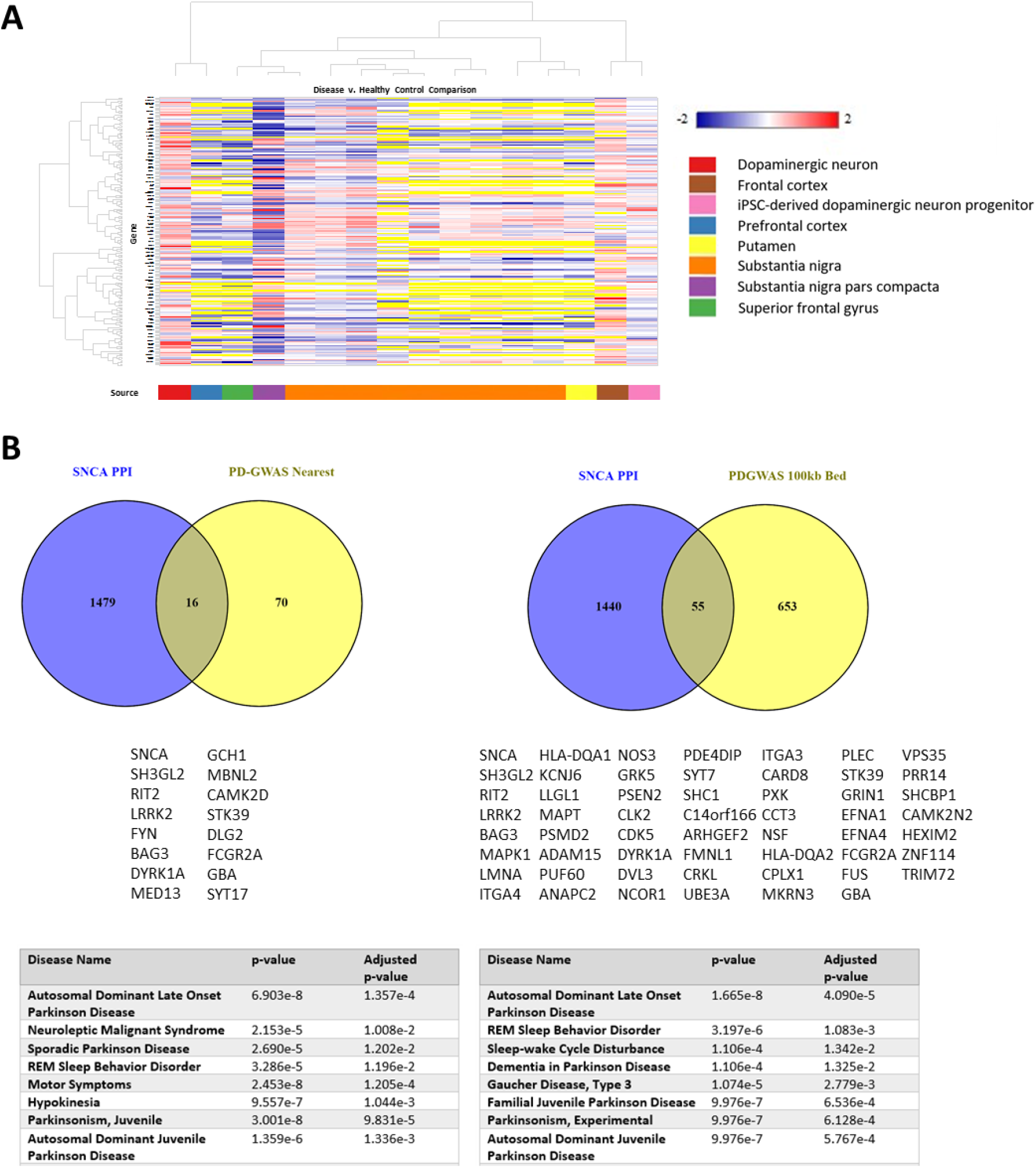
SNCA Co-expression module and PPI network genes: (A) Co-expression module genes are differentially expressed in PD Cases versus Controls in tissue samples from human cortex, putamen, and dopaminergic neuron (p-adjusted cutoff = 0.05 for differential expression in Qiagen DiseaseLand Gene Set Analysis; yellow indicates gene expression missing values). (B) PPI network gene overlaps with Nalls et al. 2019 PD-GWAS loci nearest genes and 100 kb bed genes for SNPs with p-adjusted value < 10-5 with DisGeNET set enrichment tables for genes in overlap sets.

### The identified SNCA protein-protein interactome is enriched for genes associated with PD-risk loci

We then examined which pathways within the present SNCA PPI network might be of particular relevance to synucleinopathy biology, focusing specifically on PD, for which population-level genomic analyses have previously identified a number of risk variants with high statistical significance. Among 86 unique genes mapped as the nearest genes to highly significant variants linked with PD risk in Nalls et. al, 2019, meta-analysis of genome-wide association studies (GWAS) (20), 16 were found in our SNCA PPI network (hypergeometric p = 0.0006), demonstrating significant enrichment of the SNCA PPI network for genes most closely linked with significant PD-GWAS risk loci (Fig. 7B). Furthermore, among the larger set of 708 unique genes which map to regions within ±100 kb of highly significant PD risk variants, defined as those with adjusted p-value < 10^−5^, 55 were found in the PPI network constructed from genes in the conserved SNCA co-expression module (Fig. 7B), identifying new and potentially important genes in PD pathogenesis (hypergeometric p = 0.67). Of note, enrichment of the PPI network for regionally adjacent genes is not necessarily to be expected, as this interval will contain genes with and without true contributions to PD risk. However, genes that not only have protein-protein pathway interactions with SNCA but which are also near to single nucleotide polymorphisms with highly statistically significant population-level genomic associations with PD are particularly interesting genes, as these characteristics together support a potential causal genetic contribution to disease risk.

DisGeNET queries for the overlap sets for the SNCA PPI network and the PD-GWAS nearest and ±100kb gene sets confirm enrichment of both of these sets for Parkinson’s Disease-linked genes, as well as other neurodegenerative disorders (Fig. 7B) (36). This identification of a broader set of genes with potential causal associations with PD from within the SNCA PPI network not only provides further validation of our network but also highlights the value of this newly identified network, as each node and edge represent potential targets for measuring or modulating SNCA-related activities.

### Disruption of SNCA module genes leads to elevated alpha synuclein (α-syn) levels

Genetic findings of SNCA gene duplications and triplications in familial PD strongly suggest that elevated levels of α-syn are a major contributing factor to neurodegeneration in synucleinopathies (37, 38). The SNCA co-expression network identified in the present study provides a novel platform to identify potential pathways by which different mutations might impact SNCA levels. We therefore tested whether SNCA module/PPI network genes close to PD risk variants might affect α-syn levels by knocking out a subset of these genes in human neuroblastoma cells. We selected 9 genes with the highest association to PD risk in population studies (p-value < 10^−11^) which also overlapped with the SNCA module (20). We disrupted this group of SNCA module genes singly using CRISPR knock out strategy. We observed a reduction of about 30% in specific α-syn protein staining in pooled cell lines following SNCA gene knock out in comparison to scrambled control lines. We then performed confocal microscopy imaging of fluorescently labeled α-syn for each of the SNCA module gene knock out cell lines (Figure 8A). Interestingly, there was a numerical increase in the level of synuclein staining in all but the SH3GL2 line. This increase reached statistical significance in three of the lines, STK39, GBA, and MBNL2 (Figure 8B). These findings provide experimental validation of the functional significance of the above-described SNCA PPI network.

**Figure 8.**
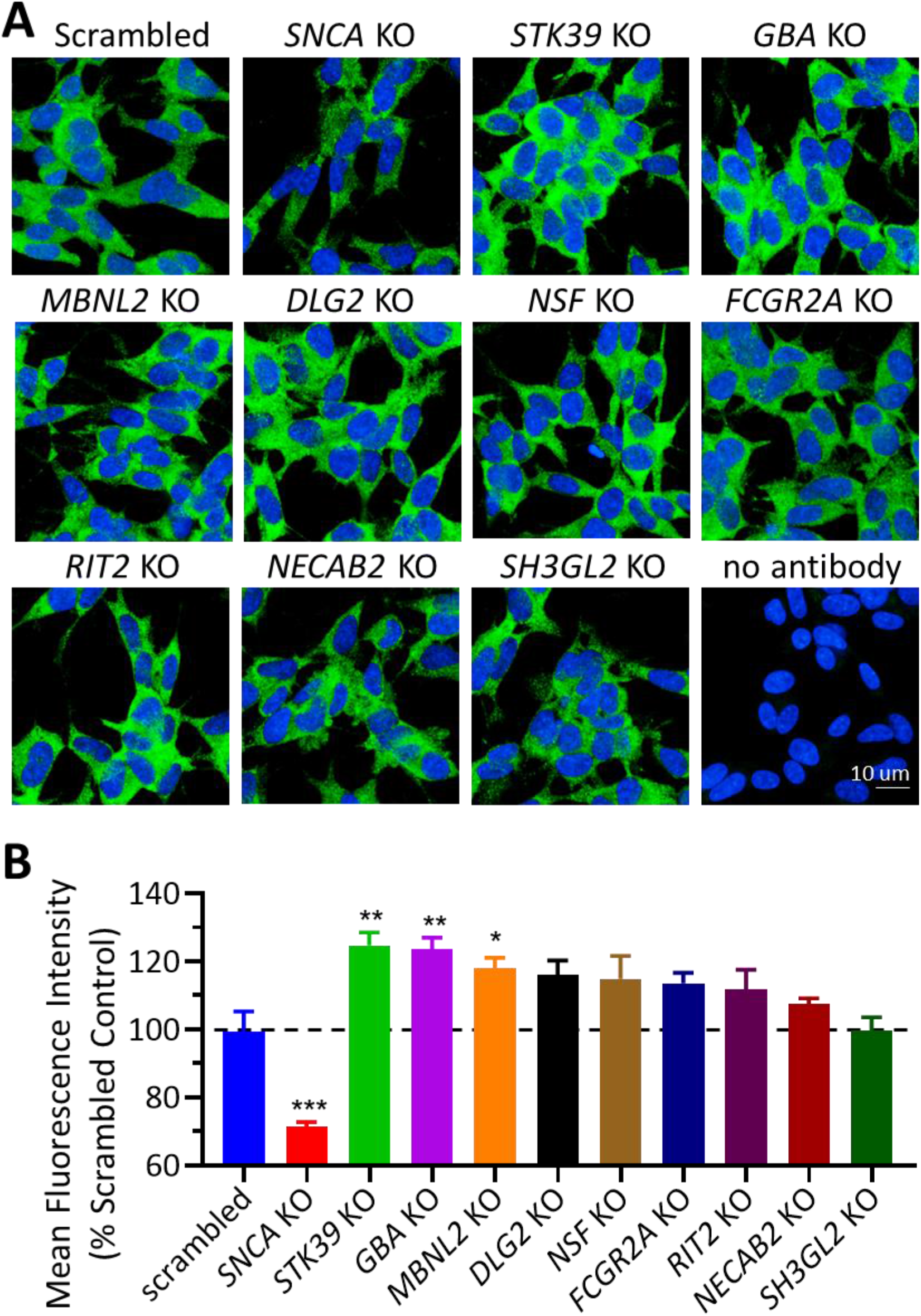
Alpha synuclein levels in CRISPR knock outs of *SNCA* module genes. Alpha synuclein protein levels are differentially regulated when *SNCA* module genes are disrupted. (A) Representative images of alpha synuclein labelled in green, and nuclei marker shown in blue in CRISPR knock outs of *SNCA* module genes. (B) Quantification of mean alpha synuclein fluorescence intensity reveals significantly elevated protein levels in *STK39*, *GBA*, and *MBNL2* knock outs in SH-SY5Y cells (*p<0.03; **p<0.003; ***p<0.001).

## DISCUSSION

The role and contributions of SNCA in different neurodegenerative disorders remains partially understood. SNCA mutations are only identified in a minority of PD and other synucleinopathy patients (39), with most synucleinopathy cases occurring in the absence of defined mutations or in the context of interacting polygenetic, epigenetic, and environmental contributions to synucleinopathy risk (6, 7). In this study, single-cell analysis reveals clear variations in SNCA expression by cell type in human ACC and MTG samples, with striking differences in SNCA expression by excitatory versus inhibitory neurons and other cell types. We also demonstrate that a number of genes implicated in synaptic biology and dopamine signaling are strongly co-expressed with SNCA in both of these cortical regions. Importantly, our finding that SNCA and its co-expressed genes are differentially expressed across neuronal cell types provokes questions of how aberrant regional or cell-type specific activity of genes linked with SNCA expression might contribute to synucleinopathy development and progression. The protein-protein interaction network constructed to connect the genes identified in our SNCA co-expression module is enriched for genes adjacent to highly significant PD risk loci found in genome-wide association studies. This network reveals new insights about the functional interactions through which a seemingly diverse set of risk variants are connected, as disease-related genes are empirically those with high propensity to interact in such network models (40).

Knock out of selected SNCA network genes further confirms the validity and functional significance of this newly characterized SNCA PPI network, with GBA, STK39, and MBNL2 identified among the knock-out cell lines tested as resulting in significantly increased SNCA protein levels relative to scrambled control. The finding that knock outs of genes within our PPI network have measurable impacts on SNCA protein levels not only supports the validity of this network but also implies that selective modulation of yet-to-be-identified SNCA PPI network pathways may also reduce SNCA levels, providing opportunities for attenuating SNCA-related disease onset or progression. To what extent SNCA expression can be modulated *in vivo* for therapeutic benefit via network interaction pathways remains to be further explored among the genes and pathways identified here. Overall, these findings provide exciting experimental evidence that disruption of SNCA module genes associated with PD risk may have functional consequences on α-syn by increasing protein levels. These findings could also further support the rationale and inform approaches for creating therapies that suppress SNCA expression with the aim of reducing total α-syn protein amounts. Such therapies might be targeted to arrest further neurodegeneration, slow down disease progression, or prevent onset of SNCA-related neuropathologies in susceptible individuals.

Examination of shortest path connections between SNCA and each of the genes found to be associated with significantly elevated SNCA protein levels in the CRISPR knock out experiments further highlights the significance and interest of the SNCA network reported in this study. In particular, STK39 (serine threonine kinase 39) was found to have the greatest effect in increasing SNCA protein expression among the genes we evaluated. STK39 is an identified PD risk gene (41, 42). Several shortest path interaction pathways connect SNCA with STK39 in our PPI network: SNCA-GABARAPL1-HSPH1-STK39; SNCA-HSP90AA1-HSPH1-STK39; and SNCA-MAP1LC3B-HSPH1-STK39. STK39 (also known as SPAK, DCHT, and PASK) phosphorylates cation-chloride coupled transporters in response to cellular stress and is an activator of the p38 MAP kinase pathway (43–45). GABARAPL1 participates in autophagic flux pathways and has been shown to be a regulator of cellular metabolism with reduction in GABARAPL1 expression shown to be associated with reduction in cellular lysosome number (46). MAP1LC3B is also an autophagy modulator (47). HSPH1 (48)and HSP90AA1(49) are both heat shock proteins participating in neuronal unfolded protein responses. These pathways identify cell-level interactions connecting SNCA with STK39 via unfolded protein response, ubiquitin, and autophagy interaction pathways. The finding that MBNL2 knock out increases cellular SNCA levels is also noteworthy and intriguing. MBNL2 (muscleblind-like 2 protein) belongs to a family of RNA-binding proteins required for terminal differentiation of neurons and other cell types (50) and has been previously implicated in the pathobiology of myotonic dystrophy as well as PD risk. Multiple shortest path connections link MBNL2 with SNCA in our network, paths with include GABARAPL1 and HSP90AA1, further implicating autophagy and unfolded protein response pathways in the etiology of PD, as these pathways are identified both in the network relationship between MBNL2 and SNCA, as well as between STK39 and SNCA as noted above.

Glucocerebrosidase (GBA) interacts directly with SNCA in our network analysis, but absent this edge, in contrast to the previous two genes, there is no other physical interaction path between GBA and SNCA in our PPI network. GBA mutations are among the most commonly identified genetic risk factors for PD (21–24), and a recent study of lysosomal storage disorder gene variants in cohort of 1156 PD patients identified the presence of at least one putative lysosomal storage disorder variant in a remarkable 56% of cases (51). GBA mutations have been reportedly identified in 3-20% of PD patients (52). GBA encodes glucocerebrosidase, a lysosomal enzyme, and reduced GBA activity has been observed among PD patients with and without GBA mutations relative to non-PD controls (53). SNCA accumulation in the setting of GBA deficiency has been proposed to occur via dysregulation of mitochondrial and autophagy pathways (54). This network analysis further identifies the essentiality of lysosomal function in preventing dysregulation of SNCA-involved pathways, providing insight into how a seemingly diverse set of pathway disruptions may converge upon synucleinopathy development, even in the absence of an identified SNCA mutation.

Another particularly interesting finding in this study was our observation of differential expression of SNCA and its co-expressed genes by neuronal cell type. in We found that SNCA and the genes co-expressed with it are most highly expressed in glutamatergic neurons. Moreover, SNCA and its co-expressed genes have functions implicated in synapse formation, synaptic function, and plasticity. These findings suggest that SNCA module genes might be playing crucial roles in regulating synaptic processes in excitatory synapses. Indeed, previous studies also discuss differential cell-type specific expression of alpha synuclein in the mouse brain (55), as well as subcellular localization of the protein to excitatory presynaptic compartments (56). Interestingly, immunolabeling of both cultured hippocampal neurons and the mouse hippocampus reveal a strong co-localization between alpha synuclein and the presynaptic vesicular glutamate transporter 1 (vGlut1), an excitatory synapse marker, suggesting a role for alpha synuclein in regulation of glutamatergic vesicles (57).

Excitatory synapses are highly plastic. They adjust their synaptic strength in response to perturbations to synaptic function throughout development, during learning and memory, disease, and aging. One way to achieve such dynamic modulation of synaptic strength is adjusting the size of the readily available, reserve, as well as the recycling pool of synaptic vesicles (58). Remarkably, alpha synuclein has been suggested to have a role in regulating recycling vesicle pool homeostasis, with one previous study showing higher levels of alpha synuclein resulted in smaller recycling pools (59). Furthermore, glutamatergic synapses, in contrast to GABAergic synapses, have a vesicle pool heterogeneity that allows for such homeostatic regulation and confers a more dynamic modulation of synaptic strength (60). As such, alpha synuclein may regulate synaptic plasticity at excitatory synapses by modulating the recycling vesicle pool size. Similarly, SNCA module genes might play pivotal roles at glutamatergic synapses in these processes by providing a scaffold for protein interactions on vesicles, regulating lipid composition and vesicle curvature, and signaling differential trafficking of biochemically diverse vesicle pools. Such roles remain speculative and await further targeted experimental validation.

In addition, more experimental data is needed to fully characterize variations in SNCA expression with perturbations in module gene expression in brain regions apart from the cortical areas profiled here. Previous work exploring transcriptional variations in the human brain has already identified regional variations in gene transcription alongside differences in cell type compositions (61). Given that each of the synucleinopathies has characteristic cellular and regional progression patterns, important questions remain regarding how cell types in each of these regions express SNCA and how these processes may be disrupted in SNCA-related pathologies. Data used for these analyses came from the MTG and ACC regions. Functions attributed to these areas include emotion and impulse regulation (ACC) (62) as well as information processing (MTG) (63). While neuropathological findings in PD initially manifest outside the cortex, cortical involvement is observed in late Braak stage disease (64, 65). Further studies are therefore necessary to examine the extent and timeline over which spatial disturbances in SNCA expression may precede the onset of various synucleinopathies. In addition, the results of our analysis highlight the particular importance of the autophagy-lysosomal pathway in the regulation of SNCA balance. How SNCA expression levels are influenced by stimuli such as environmental stress or toxic exposures, what differences in SNCA pathways may underlie cell- and region-specific vulnerabilities, and how autophagic dysregulation interacts with disease risk remain as areas for further study. For such investigations, new hypotheses and insights may be derived through consideration of possible transcriptional alterations among the SNCA-co-expressed genes identified by this analysis, as well as through new characterizations of activities along the functional pathways identified within the SNCA PPI network linking these genes.

## DATA AVAILABILITY

ACC and MTG single-nucleus data files: Allen Institute for Brain Science (https://portal.brain-map.org/atlases-and-date/rnaseq)

Seurat package software: Install from CRAN (https://satijalab.org/seurat/install.html)

WGCNA package: Installation and documentation (https://horvath.genetics.ucla.edu/html/CoexpressionNetwork/Rpackages/WGCNA/)

NetworkAnalyst online interface (https://www.networkanalyst.ca/)

Cytoscape/EnrichmentMap: gene ontology enrichment analysis applications (https://cytoscape.org/)/(apps.cytoscape.org/apps/enrichmentmap)

Enrichr: Computational systems biology interface (https://amp.pharm.mssm.edu/Enrichr/)

## SUPPLEMENTARY DATA

Supplemental Table with Guide RNA Sequences

## ACKNOWLEDGEMENT

The authors thank Deepak Rajpal for his thoughtful and constructive feedback on figure structure and content. The authors thank Meaghan Cogswell for her insightful feedback and review of this project.

---

**Table S1.**
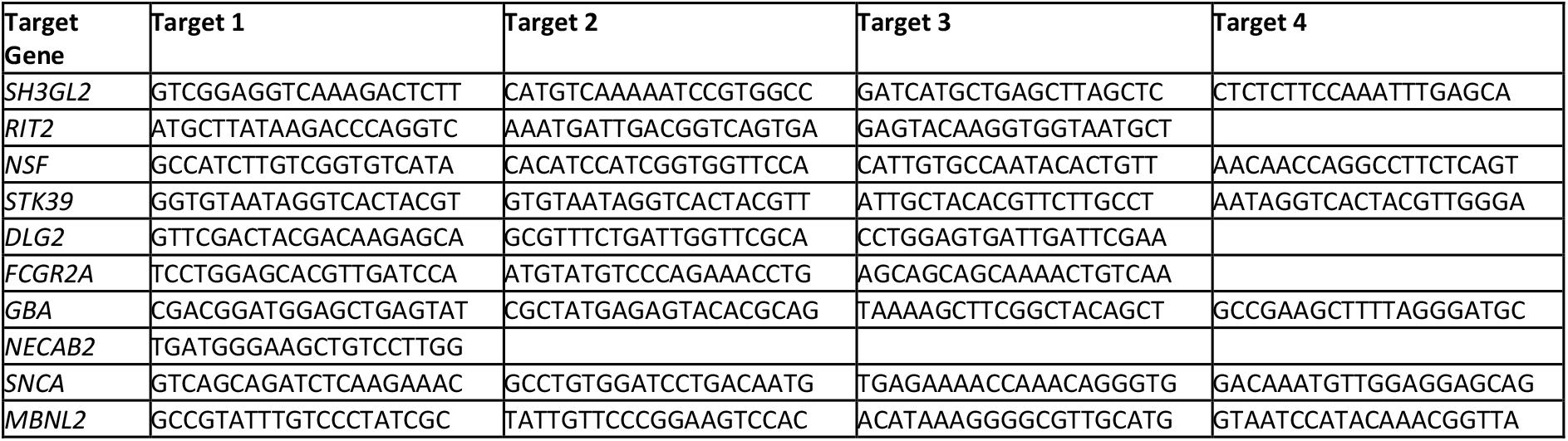
Supplemental table with guide RNA Sequences.

